# Carriage of three plasmids in a single human clinical isolate of *Clostridioides difficile*

**DOI:** 10.1101/2022.07.10.499461

**Authors:** Anna M. Roseboom, Quinten R. Ducarmon, B.V.H. Hornung, C. Harmanus, M.J.T. Crobach, Ed J. Kuijper, Rolf H.A.M. Vossen, Susan L. Kloet, Wiep Klaas Smits

**Affiliations:** Department of Medical Microbiology and Leiden University Center of Infectious Diseases (LU-CID), Leiden University Medical Center, Leiden, The Netherlands; Department of Human Genetics, Leiden Genome Technology Center, Leiden University Medical Center, Leiden, The Netherlands; Centre for Microbial Cell Biology, Leiden, The Netherlands

**Keywords:** Complete genome sequence, plasmid compatibility, *C. difficile* carriage

## Abstract

A subset of clinical isolates of *Clostridioides difficile* contains one or more plasmids and these plasmids can harbor virulence and antimicrobial resistance determinants. Despite their potential importance, *C. difficile* plasmids remain poorly characterized. Here, we describe a human clinical isolate that carries three plasmids from three different plasmid families that are therefore compatible. For two of these, we identify a region capable of sustaining plasmid replication in *C. difficile*. Together, our data advance our understanding of *C. difficile* plasmid biology.

**Highlights:**

The complete circular genome sequence is provided for a *C. difficile* isolate harboring three plasmids
These three plasmids (pJMR5-1, pJMR5-4 and pJRM5-W) are therefore compatible in a single strain
Sequence analysis suggest a modular nature of plasmid families to which the pJMR-plasmids belong
A functional replicon was cloned from pJMR5-1 (pCD-ECE1 family) and pJMR5-W (pCD-WTSI1 family) and plasmids carrying this replicon are compatible with plasmid pCD630

## (1) Introduction

The Gram-positive anaerobic spore-forming bacterium *Clostridioides difficile* is responsible for healthcare-associated and community-acquired infectious diarrhea with potentially fatal consequences [1]. The symptoms of a *C. difficile* infection (CDI) are related to the expression of one or more toxins, but virulence of this bacterium is multifactorial [2].

In many Gram-positive pathogens, virulence factors are encoded on plasmids [3, 4]. For *C. difficile*, sporadic reports indicate that toxins and resistance determinants can be carried on extrachromosomal elements [5–7]. It is estimated that ~10-70% of *C. difficile* isolates carry one or more plasmids, but information on plasmid functions is sparse [8]. In particular, very few replicons have been identified and almost no experimental evidence on plasmid compatibility is available. Regions sufficient for replication in *C. difficile* have been cloned from the plasmids pCD6 and pCD-METRO from strains CD6 and IB136, respectively [5, 9]. An *in silico* analysis of publicly available sequence data suggests that specific families of plasmids may co-exist [8, 10], but these predictions have not been validated experimentally.

Here, we show carriage of plasmids from three different plasmid families in a single isolate of *C. difficile* derived from a human patient and report on the identification of a region sufficient for plasmid maintenance for two of these plasmids. Together, these data significantly advance our understanding of *C. difficile* plasmid biology.

## (2) Materials and methods

### (2.1) Bacterial strain and growth conditions

All bacterial strains are listed in **Table 1**. The isolates used in this study are derived from the collection of isolates at the Dutch national Reference Laboratory (NRL) for *C. difficile*, which is housed at the Leiden University Medical Center. Strains JMR1, JMR2, and JMR5 have been isolated from one or more patients at the same healthcare facility in the context of diagnostics. These strains were collected over a period of five months and were sent to the NRL for sentinel surveillance. Strain JMR7 was collected in the context of a multicenter study aimed at determining the value of *C. difficile* colonization screening at hospital admission in an endemic setting [11].

**Table 1.** Strains.

| Strain | Description | Reference |
| --- | --- | --- |
| ***Clostridioides difficile*** |  |  |
| JMR1 | Human clinical isolate; RT081. Contains three plasmids. | This study |
| JMR2 | Human clinical isolate; RT081. Contains three plasmids. | This study |
| JMR5 | Human clinical isolate; RT081. Contains pJMR5-1, pJMR5-4 and pJMR5-W plasmids. | This study |
| JMR7 | Human clinical isolate; RT081. Contains three plasmids. | This study |
| 630Δerm | Laboratory strain; RT012. | [24, 31] |
| JIR8094 | Laboratory strain; RT012. | [36] |
| DSMZ 28645 | Laboratory strain; RT012. | [35] |
| ***Escherichia coli*** |  |  |
| AP24 | DH5α x pAP24 (pRPF185-derived vector containing sluc<sup>opt</sup> and pCD6 replicon, catP) | [23] |
| DH5α | *fhuA2 lac(del)U169 phoA glnV44 Φ80' lacZ(del)M15 gyrA96 recA1 relA1 endA1 thi-1 hsdR17* | Laboratory stock |
| JMR20 | DH5α x pJMR20 (pAP24-derived vector containing pJMR5-1 ORF8 and flanking intergenic regions) | This study |
| JMR25 | DH5α x pJMR25 (pAP24-derived vector containing pJMR5-1 ORF7-ORF10 and flanking intergenic regions) | This study |
| JMR28 | DH5α x pJMR28 (pAP24-derived vector containing pJMR5-W ORF1 and flanking intergenic regions) | This study |
| JMR33 | DH5α x pJMR33 (pAP24-derived vector containing pJMR5-4 ORF5 and flanking intergenic regions) | This study |
| JMR46 | DH5α x pJMR46 (pAP24-derived vector containing pJMR5-4 ORF1-5 and flanking intergenic regions) | This study |
| JMR51 | DH5α x pJMR51 (pAP24-derived vector containing pJMR5-1 ORF8-10 and flanking intergenic regions) | This study |
| JMR52 | DH5α x pJMR52 (pAP24-derived vector containing pJMR5-1 ORF7-8 and flanking intergenic regions) | This study |
| JMR57 | DH5α x pJMR57 (pAP24-derived vector containing pJMR5-W ORF1-4 and flanking intergenic regions) | This study |

Isolates were routinely cultured on TSS plates (Tryptic Soy Agar with 5% sheep blood, bioMérieux, The Netherlands) or CLO plates (selective *C. difficile* medium containing cefoxitin, amphotericin B and cycloserine, bioMérieux, The Netherlands). Capillary electrophoresis PCR ribotyping was performed at the Dutch National Reference Laboratory for *C. difficile*, according to standard procedures [12] and further characterized using a multiplex PCR targeting the 16S rRNA gene, *gluD*, and the genes encoding the large clostrididal toxins and binary toxin [13].

Laboratory strains of *C. difficile* were cultured anaerobically at 37°C in liquid BHIY (1.5 % w/v Brain Heart Infusion [Oxoid], 0.5% w/v yeast extract [Sigma]) or on BHIY agar plates (BHIY, 1.5% agar w/v), supplemented with *C. difficile* selective Supplement (CDSS [Oxoid]) and 25 μg/mL thiamphenicol when appropriate, in a Don Whitley VA-1000 workstation (10% CO2, 10% H2 and 80% N2 atmosphere). *E. coli* strains were cultivated aerobically at 37°C, 200 rpm in Luria-Bertani (LB) broth or on LB agar plates, supplemented with 50 μg/mL ampicillin, 25 μg/mL chloramphenicol and/or 50 μg/mL kanamycin when required. Stocks were made in 15% w/v glycerol and stored at −80°C.

### (2.2) DNA isolation and whole genome sequencing of *C. difficile*

Initial analysis of the draft genome of strain JMR5, including the tentative identification and annotation of extrachromosomal elements, was performed based on short-read Illumina sequence data (ERS2564723 from PRJEB25045) as described [10, 11]. The complete genome was generated based on long read sequencing on the Pacific Biosciences (PacBio) Sequel platform. To prepare high molecular weight total DNA, cells from 5 mL of overnight culture were pelleted and processed using the Qiagen Genomic-tip 100/g, according to the manufacturer’s instructions. PacBio sequencing libraries were generated according to the manufacturer’s multiplexed microbial library preparation protocol, part number 101-696-100, version 7, July 2020 release using the SMRTbell Express Template Prep Kit v2.0 with the following modifications: genomic DNA was sheared using Speed 34 on the Megaruptor 3 (Diagenode) and an additional size selection step of 6-50kb fragments on the Blue Pippin (Sage Science) was included for the final SMRT bell library. The libraries were sequenced on a Sequel II platform (Pacific Biosciences) using the Sequel II Binding Kit v2.0, Sequencing Primer v4, Sequencing Kit v2.0 and a 30hr movie time.

### (2.3) Data analysis and visualization

Raw PacBio sequence data was assembled using Flye (v2.9)[14] and the start position was fixed using Circlator and the “fixstart” parameter [15]. Assembly quality was subsequently inspected using QUAST [16] and assembly completeness using BUSCO (v5.3.2, dataset clostridia_odb10 creation date 2020-03-06)[17]. Prokka (v1.14.6) was used for rapid genome annotation and to obtain protein sequences, with the “–kingdom Bacteria”, “--genus Clostridioides” and “–species difficile” parameters [18]. Multi-locus sequence typing (MLST) was performed using mlst (v2.19.0) with the PubMLST *C. difficile* database updated to October 21^st^ 2021 [19]. Alignments of plasmid families were visualized using clinker [20]. Amrfinderplus (v3.1.23)[21] with the database version of December 21^st^ 2021 was run to identify acquired antimicrobial resistance (AMR) genes, genes with point mutations conferring resistance to antimicrobials (specifically *gyrA, gyrB, murG, rpoB, rpoC* and *23S*) and virulence factors (including the *C. difficile* toxin genes) with the following options: “-n” “--organism Clostridioides_difficile”, “—plus”. The MobileElementFinder webserver was used to detect potential mobile elements with default settings.[22].

### (2.4) Data and code availability

Raw sequence data generated for this study is available at the European Nucleotide Archive under BioProject PRJEB53950. All bioinformatic tools used for the analyses are freely available through the references provided. Plasmid sequences have also been deposited independently as ON887052 (pJMR5-1), ON887053 (pJMR5-4) and ON887054 (pJMR5-W) in GenBank. The complete genome sequence of JMR5 is GCA_944989955.1 (assembly)/ERS12289077 (sample). Previously generated whole genome sequences [11] are available under BioProject number PRJEB25045. Sequences for pCD-ECE1 (LR594544.1), pCD-ECE4 (LR594545.1), pCD630 (AM180356.2) and pCD-WTSI1 (MG019959.1) were retrieved from GenBank.

### (2.5) Confirmation of extracellular nature of the plasmids

In order to confirm the extrachromosomal nature of the plasmids, a PlasmidSafe DNase (PSD; Epicentre) experiment was performed as described previously [5]. In short, total genomic DNA was isolated and incubated in the presence or absence of PSD for 24h. Chromosomal DNA is fragmented and susceptible for degradation by PSD in this procedure, whereas small circular double stranded DNA (plasmids) is not. After incubation, the remaining DNA was purified and amplified with primers specific for loci on the chromosome (oWKS-1070/oWKS-1071), pJMR5-1 (oAR-1/oAR-2), pJMR5-4 (oAR-3/oAR-4) and pJMR5-W (oAR-5/oAR-6) using MyTaq DNA polymerase (Meridan) according to the instructions of the manufacturer. The sequences of all oligonucleotides are listed in **Table 2.**

**Table 2.**
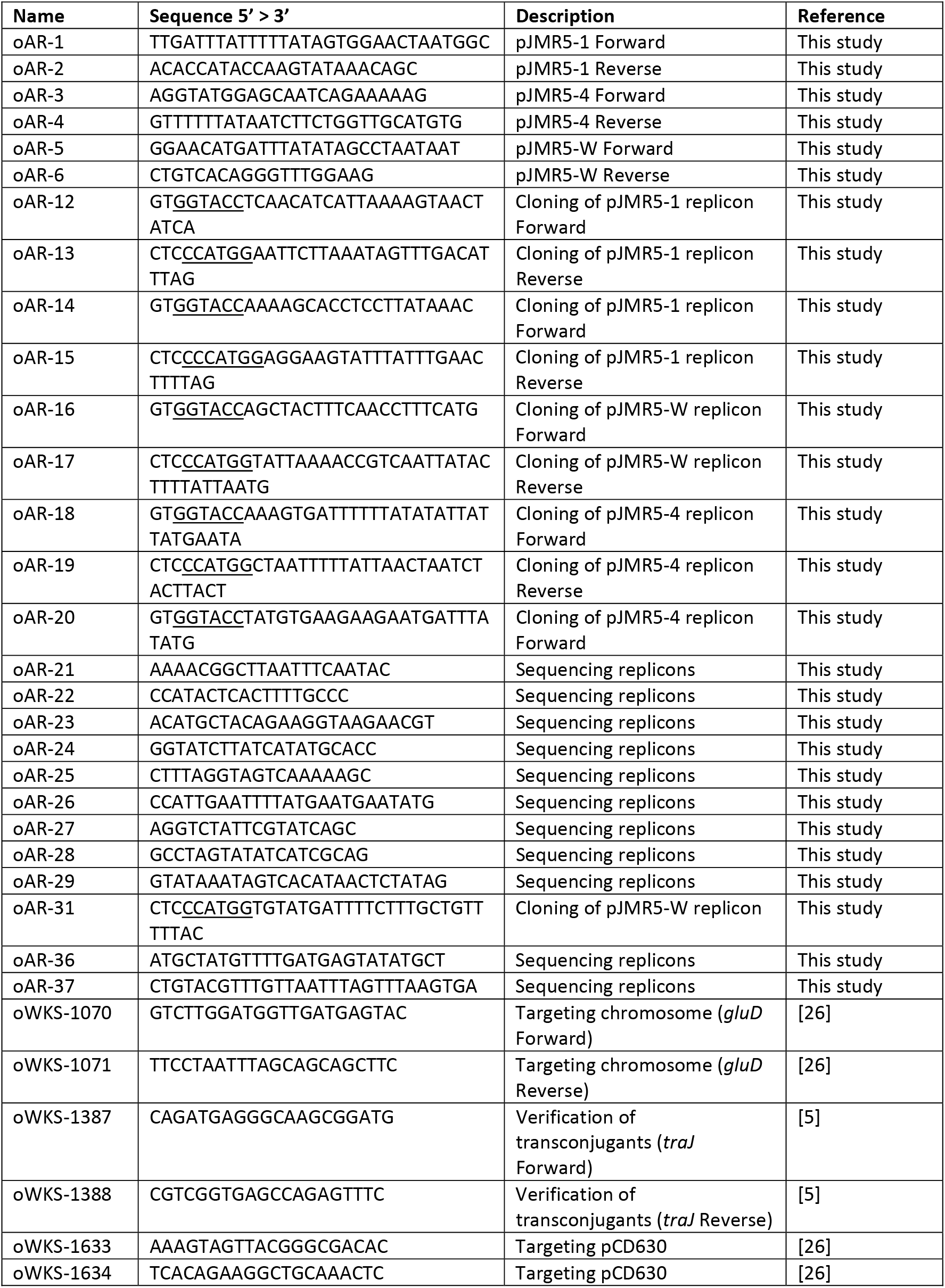
Oligonucleotides used in this study. Relevant restriction sites are underlined.

### (2.6) Molecular biology procedures

All vectors were constructed with restriction/ligation cloning using a linearized pAP24-based vector [23] as the backbone and a DNA fragment of interest as insert. Plasmids were isolated from 1 mL overnight culture using NucleoSpin Plasmid Easypure columns (Macherey-Nagel) from the Plasmid Easypure Kit (Nucleospin).

Putative replicon fragments were obtained through amplification PCR using high fidelity enzyme Q5 DNA polymerase (NEB) and primers including a KpnI or NcoI restriction site (**Table 2**). After digestion with KpnI/NcoI, the amplicons were ligated into similarly digested pAP24-backbone [23], as described previously [5], and transformed to *E.coli* DH5α with selection on chloramphenicol-containing LB agar plates. All constructs were verified by Sanger sequencing using primers listed in **Table 2**.

For conjugation, *E.coli* CA434 competent cells [9] were transformed using purified plasmid DNA and transformants were selected on LB agar supplemented with chloramphenicol and kanamycin. Conjugative transfer of the plasmids from CA434 to *C. difficile 630Δerm* [24] was performed according to standard procedures [9]. The transconjugants, selected on thiamphenicol containing media, were screened with PCR using MyTaq DNA polymerase (Meridan) directly on colony material or on purified DNA. DNA extractions were performed using the DNeasy Blood and Tissue (Qiagen) kit after enzymatic cell lysis as prescribed by the manufacturer. To confirm identity of the transconjugants, the presence of *gluD* (on the chromosome) and *traJ* (on the plasmid backbone) was verified in a PCR using primers oWKS-1070/oWKS-1071 and oWKS-1387/oWKS-1388, respectively (**Table 2**). The presence of the pCD630 plasmid in the DNA samples of the TCs was confirmed by performing a PCR using primers oWKS-1633 and oWKS-1634 (**Table 2**).

## (3) Results and Discussion

### (3.1) Characterization of the plasmid-containing isolates

Three of the four clinical isolates (JMR1, JMR2 and JMR5) have been isolated from a single patient in a single healthcare facility as part of the ongoing activities of the Dutch National Reference Laboratory for *C. difficile*. Medical records show that the patient initially experienced mild symptoms, but eventually succumbed to the infection after several more CDI episodes. The fourth isolate (JMR7) was collected as part of a study looking at *C. difficile* colonization at hospital admission [11]. As the patient did not give full consent during this study [11], only limited metadata is available for this isolate. We note, however, that JMR5 and JMR7 were collected in the same time period and place, from patients of the same age and gender. Therefore, it is likely that JMR1, JMR2, JMR5, and JMR7 are all longitudinal samples from the same patient suffering from recurrent CDI over a period of at least 4 months. If the samples are indeed derived from the same patient, we would expect all *C. difficile* isolates to be similar in terms of PCR ribotype and toxin profile.

Therefore, we performed capillary PCR ribotyping and a multiplex PCR targeting the toxin genes. This data showed that all JMR isolates belong to PCR ribotype 081; this PCR ribotype is reported to be part of the phylogenetic clade 1 of non-epidemic isolates and is not among the most commonly found PCR ribotypes in Europe [25]. In agreement with the ribotyping result, the multiplex PCR shows that all isolates are toxin A and toxin B positive, but negative for binary toxin.

Additionally, we extracted the relevant core genome MLST information (cgMLST) from the previously generated Illumina sequences (GenBank PRJEB25045)[11]. This showed that all isolates belong to the same cluster type, with 0 allele differences. Together, these data support the notion that they are derived from a single persistently infected individual.

### (3.2) Confirmation of the extrachromosomal nature of pJMR5-1, pJMR5-2 and pJMR5-3

Isolate JMR7 was subjected to short-read whole genome sequencing, as part of the aforementioned study. During the analyses, carriage of extrachromosomal elements was predicted, according to in-house established methods [10]. The predictions indicate possible extrachromosomal elements belonging to the pCD-ECE1, pCD-ECE4 and pCD-WTSI1 families, of 6.5, 15.4 and 21.9 kb size, respectively.

We extracted the contigs corresponding to the predicted extrachromosomal elements, and circularized them after removal of the terminal repeats. To confirm the presence and the circular dsDNA nature of the elements *in vivo*, we performed a PlasmidSafe DNase (PSD) analysis using primers specifically directed at regions of each putative plasmid [26]. The analysis showed that a product of the expected size was obtained in PCRs targeting each of the putative plasmids, even for PSD-treated samples. In contrast, a PCR product for the chromosomal locus *gluD* was only obtained in the samples that were not treated with PSD (**Figure 1**). We note a decrease in the strength of the signal on gel for the pCD-WTSI family plasmid upon treatment with PSD; as this is the largest of the predicted plasmid, we attribute this to partial fragmentation of the dsDNA during the DNA isolation.

**Figure 1.**
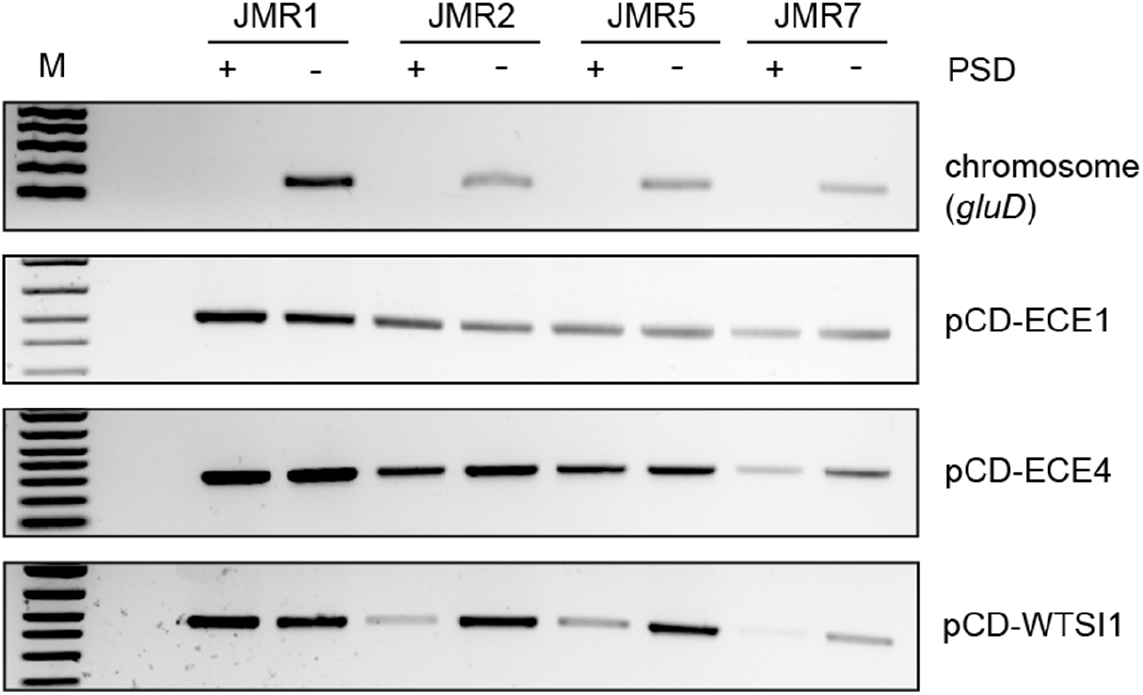
JMR strains contain three plasmids. Total genomic DNA was incubated in the absence (-) or presence (+) of PlasmidSafe DNase (PSD) and subsequently amplified with primers specific for the indicated DNA (pCD-ECE1 family: oAR1/2, pCD-ECE4 family: oAR3/4, pCD-WTSI1 family: oAR4/5; see **Table 2** and Materials and Methods).

These data indicate that all JMR strains contain circular dsDNA that is extrachromosomal, consistent with the presence of a plasmid.

### (3.3) Complete genome sequence of *C. difficile* JMR5

Next, we determined the complete genome sequence of JMR5, as a representative of the group of JMR strains, using single molecule real time sequencing. Assembly of the reads of the Pacific Bioscience Sequel platform yielded a single circular chromosome of 4.3 Mb and 3 smaller replicons (**Supplemental Figure 1**). The size and sequence of the latter matched the predicted circular elements discussed above, and are hereafter referred to as pJMR5-1 (for **p**lasmid from **JMR5** from the pCD-ECE**1** family), pJMR5-4 (for **p**lasmid from **JMR5** from the pCD-ECE**4** family) and pJMR5-W (for **p**lasmid from **JMR5** from the pCD-**W**TSI1 family). Initially, we recovered only pJMR-4 and pJMR5-W in the *de novo* assembly, possibly as a result of size selection during the library preparation (see Materials and Methods). However, manual inspection of the data revealed a significant number of reads matching pJMR5-1 and these reads fully matched the predicted sequence; indeed, altering the default settings of Flye [14] allowed automatic recovery of the pJMR5-1 plasmid, though in this case the plasmid contained a duplication (a well-known artefact of long-read sequencing of small plasmids) [15]. A split library preparation (<10kb and 10+ kb) may improve automatic recovery plasmids of <10kb in a de novo assembly using Flye.

The genome of JMR5 is 4.321.867 bp and has an average [G+C]-content of 29%, in line with other *C. difficile* genomes. An automated annotation identifies 3937 ORFs, 35 rRNA genes, 90 tRNA genes and a single tmRNA. Several AMR genes were identified by AMRfinderplus [21] on the JMR5 chromosome, namely bla-CDD1, erm(B), the vanGCd operon and a T82I point mutation in *gyrA*, associated with fluoroquinolone resistance. One AMR gene, tet(M), was present on pJMR5-W. No mobile elements were identified by MobileElementFinder [22].

We used PubMLST [19] to predict the sequence type of JMR5. The isolate was identified as belonging to ST9, consistent with previous studies that place the RT081 reference strain from the Leeds-Leiden collection in the same sequence type [27, 28].

### (3.4) Comparison of the identified plasmids with reference plasmids

The tentative assignment of the three plasmids to the pCD-ECE1, pCD-ECE4 and pCD-WTSI1 family of plasmids is based on regions of similarity with the reference sequences for these plasmids as described [10]. To gain more insight in the relatedness of the plasmids, we generated and visualized nucleotide alignments using clinker [20]. From these analyses, multiple observations were made.

First, we note that the ORF7-ORF8 region of pCD-ECE1 appears not to be conserved in pJMR5-1 (**Figure 2A**). This was unexpected, as ORF8 of pCD-ECE1 was hypothesized to encode a replication-associated protein based on limited homology to viral REP proteins [10]. If this region of the plasmid is involved in replication, the corresponding region in pJMR5-1 should also encode a replication protein. Indeed, analysis of the predicted amino acid sequence of the pJMR5-1 protein using the PHYRE2 Protein Fold Recognition Server [29] indicates that it likely is a rolling circle replication initiator protein (PDB 4CIJ; confidence 98.8%; identity; 13%).

**Figure 2.**
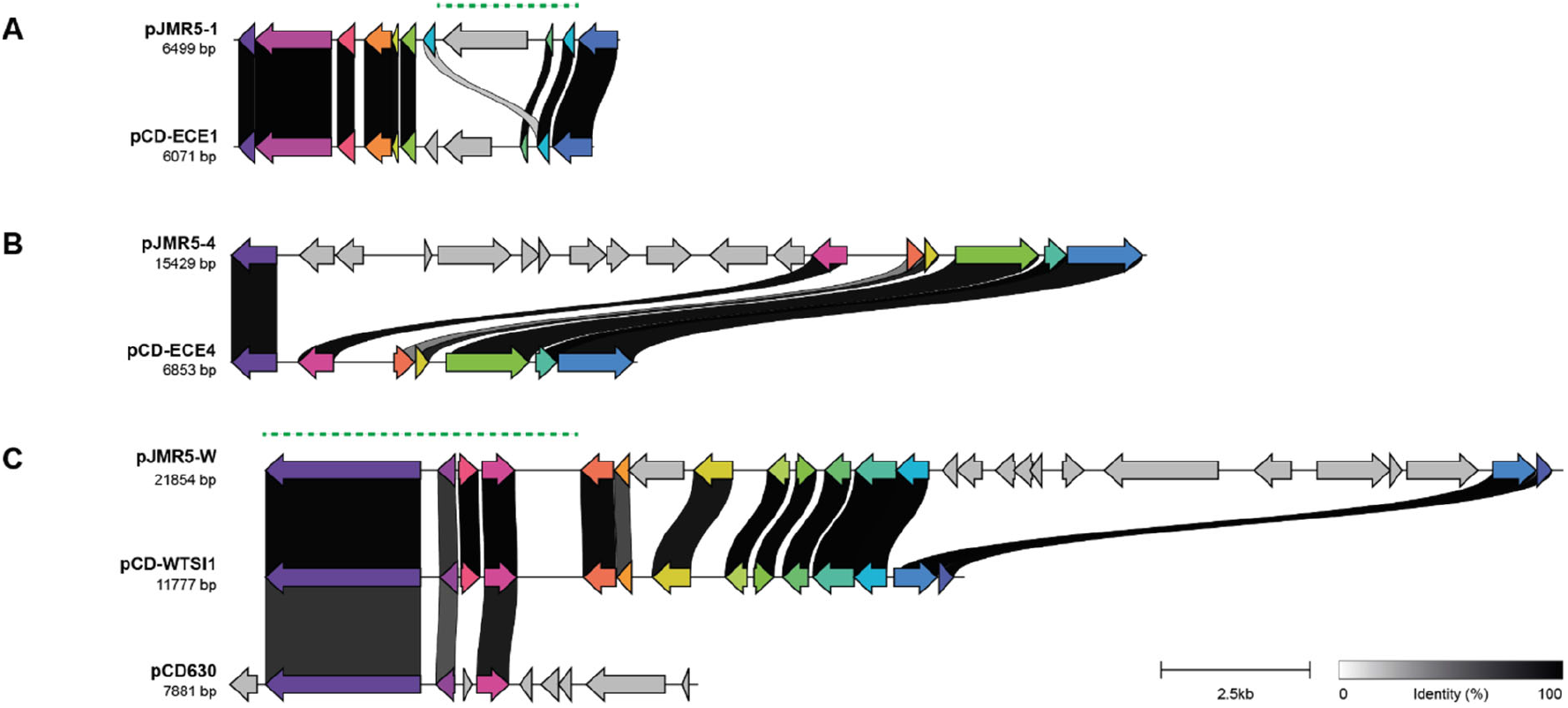
Comparison of pJMR5 plasmids with reference plasmids. Within panels, colored arrows indicate similar genes; links are drawn between similar genes on neighboring clusters and are shaded based on sequence identity (0% white, 100% black, identity threshold for visualization 0.30). Dashed green lines indicate regions capable of sustaining replication of a shuttle vector in *C. difficile*. **A**. Comparison of pJMR5-1 (ON887052) with pCD-ECE1 (LR594544.1). **B**. Comparison of pJMR5-4 (ON887053) with pCD-ECE4 (LR594545.1). **C**. Comparison of pJMR5-W (ON887054) with pCD630 (AM180356.2) and pCD-WTSI1 (MG019959.1). Image is based on a visualization with clinker [20].

Second, the pJMR5-4 plasmid appears to contain a 11-gene insertion in comparison to the pCD-ECE4 reference sequence (**Figure 2B**), consistent with a previously reported large size-diversity within this family [10]. Automated annotation of gene function (see Materials and methods) suggests that this region may include a methylation sensitive restriction modification system as ORF12 of pJMR5-4 encodes a putative type II endonuclease, and ORF11 encodes a DNA (cytosine-5-)-methyltransferase.

Third, pJMR5-W is substantially larger than other members of the pCD630/pCD-WTSI1 family (**Figure 2C**)[26]. Plasmids pCD630 and pCD-WTSI1 share a conserved region that encodes a helicase protein, believed to be important for replication, but differ in their accessory genes [26]. pJMR5-W also contains the conserved region but it appears that pJMR5-W contains two insertions compared to pCD-WTSI1; i) a single hypothetical gene that is also found at this site for other pCD-WTSI family plasmids [26] and ii) an 11-gene cluster suggestive of a mobile element (e.g. including a gene encoding an Tn*916*-like excisionase [ORF23] and an integrase [ORF24]). Interestingly, PHYRE2 [29] also predicts multiple regulator proteins (ORF14, ORF15, ORF19, ORF21), a putative rolling circle replication initiator protein (PDB 4CIJ, confidence 100, 28% identity), and a putative tetracycline resistance protein (ORF20)(PDB 3J25, confidence 100, 97% identity) in the island. Tn*916*-like elements are common vehicles for tetracycline resistance genes [30].

### (3.5) A functional replicon from pJMR5-1 and pJMR5-W

Based on the sequence analysis, we expected ORF8 of pJMR5-1 to be required for replication of the plasmid in *C. difficile*. To assess which region is sufficient for replication, we replaced the pCD6 replicon from an established *E.coli-C. difficile* shuttle vector [23] with different, but overlapping, regions of the pJMR5-1 plasmid. The obtained constructs were conjugated into the *C. difficile* laboratory strain *630Δerm* [24, 31]; when transconjugants were obtained this was taken as a sign of a functional replicon, whereas repeated failure to obtain transconjugants suggested that a critical determinant was missing. A region encompassing ORF7-ORF10 sustained plasmid replication, as did a region encompassing ORF8-ORF10 as transconjugants were readily obtained when conjugating plasmids pJMR25 and pJMR51 (**Table 3**). In contrast, a region encompassing only ORF8, or ORF7-ORF8 did not, as we failed to obtain transconjugants with plasmid pJMR20 and pJMR52 (**Table 3**). We conclude that the ORF8-ORF10 is sufficient to allow the plasmid to replicate (**Figure 2A**). Furthermore, preliminary data suggests that ORF10, encoding an Arc-type ribbon-helix-helix protein is essential for this function (data not shown).

**Table 3.** Results of conjugations.

| Origin of putative replicon | Plasmid | Recipient strain | pCD630 in recipient (yes/no) | Transconjugants obtained (yes/no) |
| --- | --- | --- | --- | --- |
| pJMR5-1 | pJMR20 | 630Δ*erm* | Yes | No |
| | pJMR25 | 630Δ*erm* | Yes | Yes |
| | pJMR51 | 630Δ*erm* | Yes | Yes |
| | pJMR52 | 630Δ*erm* | Yes | No |
| pJMR5-4 | pJMR33 | 630Δ*erm* | Yes | Yes |
| | pJMR46 | 630Δ*erm* | Yes | No |
| pJMR5-W | pJMR28 | 630Δ*erm* | Yes | No |
|  |  | JIR8094 | No | No |
|  |  | DSMZ 28645 | No | No |
| | pJMR57 | 630Δ*erm* | Yes | Yes |
|  |  | JIR8094 | No | Yes |
|  |  | DSMZ 28645 | No | Yes |

In 2019, analysis of the ORFs on the pCD-ECE4 plasmid failed to predict a replicon region in plasmids from this family [10]. Screening the ORFs of pJMR5-4 using PHYRE2 indicated that ORF5 likely encodes a protein of which the core domain shows homology to the archaeo-eukaryotic primase (AEP) domain of an archael primase protein (PDB 1V33; confidence 96.8, 32% identity)[32]. Primase-polymerases (PrimPols) are archaeo-eukaryotic primases (AEP) with both primase activity and DNA polymerase activity [33]. The AEP supergroup PrimPol-PV1 is widespread in diverse bacterial and archaeal plasmids and likely plays a crucial role in plasmid replication [34]. However, a region encompassing this protein did not sustain plasmid replication in *C. difficile* (**Table 3**). A larger region, additionally encompassing ORF1-ORF5 and an area of high repeat-density, similarly failed to sustain replication (**Table 3**). We conclude that this region does not contain a functional replicon. It is likely that one or more of the hypothetical genes in the conserved part of the pCD-ECE4 family of plasmids encodes the replication function.

The plasmid pJMR5-W contains the helicase-containing module that defines the pCD630/pCD-WTSI family of plasmids [26]. We cloned two different fragments of the highly conserved region into a vector as potential replicons; pJMR57 carries the complete conserved region as putative replicon, and pJMR28 includes only ORF1, encoding the putative helicase protein. We considered that the pJMR5-W replicon might not be compatible with the replicon of pCD630 due to high similarity (**Figure 3B**). Plasmid pCD630 is present in our recipient strain *630Δerm* [24] so two additional recipient strains were included; DSMZ 28645 (a 630Δ*erm* strain from the Leibniz Institute DSMZ collection)[35] and JIR8094 (also known as 630E)[36] both of which do not contain the pCD630 plasmid [26]. Conjugation of the three strains with pJMR28 (containing only the ORF9 fragment) repeatedly did not result in any growth on the selective plates, while conjugation with pJMR57 (containing the complete conserved region) did yield viable transconjugants carrying the vector. Therefore, we conclude that the conserved region characterizing all pCD630/pCD-WTSI plasmids comprises a functional replicon, whereas a fragment encoding only the helicase is not sufficient for plasmid maintenance. The fact that plasmid pJMR57 was successfully introduced into a recipient strain carrying pCD630 indicates that pCD630 and pCD-WTSI1 are compatible plasmids, at least for the duration of the experiment. This was confirmed by a PCR specifically targeting the pCD630 sequence on purified DNA from the obtained transconjugants (data not shown). Our experimental data is consistent with the *in silico* prediction that pCD-WTSI family plasmids appear compatible with most other plasmids, including others from the pCD-WTSI1 family (i.e. the replication function does not confer incompatibility)[8]. Of note, our results also show that the putative rolling circle replication protein of pJMR5-W (ORF22) – that is not present in pCD-WTSI1 or pCD630 - is not strictly required for replication, despite its structural similarity to a plasmid replication protein. However, we have not yet established whether a region of the plasmid that incorporates this protein might also be able to sustain replication so we cannot rule out such a function at this time.

## (4) Conclusions

In this study we demonstrated carriage of three different plasmids in a clinical isolate obtained from a symptomatic patient suffering from rCDI. Our results suggest that for *C. difficile* i) non-conserved regions of putative plasmid families can be involved in plasmid replication (as exemplified by pJMR5-1), ii) plasmid replication proteins may exist that defy current *in silico* predictions of protein structure and function (pJMR5-4) and iii) the conserved module of a wide-spread plasmid family can sustain plasmid replication (pJMR5-W). Importantly, epidemiological evidence suggests that the pJMR5-1, pJMR5-4 and pJMR5-W plasmids are retained during a persistent infection over a period of months.

## Acknowledgements

The authors would like to acknowledge B. Nibbering for isolating the total genomic DNA for PacBio sequencing, S. Nooij for assistance with PacBio-data analysis and I. Sidorov for the local implementation of clinker at the LUMC.

## Funding

This research did not receive any specific grant from funding agencies in the public, commercial, or not-for-profit sectors.

## Declarations of interest

none

## Author contributions

AR: investigation, formal analysis; QRD: formal analysis, writing – original draft, visualization; BVHH: conceptualization, methodology, formal analysis; CH: investigation; MJTC: formal analysis; EJK: supervision; RHAMV: investigation, formal analysis; SLK: project administration, supervision, resources; WKS: conceptualization, formal analysis, visualization, project administration, resources, writing – original draft. All authors contributed to review & editing.

## Notes

### Competing Interest Statement

The authors have declared no competing interest.

